# The Nigro-Thalamic Projection contributes to the Control of Action Initiation Timing

**DOI:** 10.1101/2020.01.12.903500

**Authors:** Julien Catanese, Dieter Jaeger

## Abstract

The nigro-thalamic pathway is one of the main outputs from the basal ganglia, known to be involved in action selection, learning motor skills, and/or control the vigor of actions. However, the specific function of the nigral input to the motor thalamus remains unclear. Using a combination of in vivo electrophysiological recordings from motor thalamic neurons and optogenetic activation of nigral inputs in behaving head-fixed mice, we determined that this pathway is primarily involved in the proper timing of the release of goal directed actions. At the behavioral level, we were able to reduce the amount of impulsive licking by activating thalamic terminals. We describe changes in thalamic neuronal activity explaining this effect and propose a parsimonious model to account for our observations. These results provide new insight in the circuitry for timing control, a critical part of motor planning, and reveal a potential new target to modulate impulsivity.

## INTRODUCTION

The basal ganglia (BG) have been implicated in action selection (Donahue and Kreitzer, 2015; Mink, 1996), motor learning (Graybiel, 2008), in the control of movement vigor (Turner and Desmurget, 2010), and timing of movement initiation (Buhusi and Meck, 2005). The classic conceptual model of basal ganglia circuitry posits that these functions are conveyed through a cortico-basal ganglia thalamocortical loop (Alexander and Crutcher, 1990). A critical link in the classic loop pathway is the motor thalamus, where in the rodent, inhibitory BG output from the Substantia nigra reticulata (SNr) and entopeduncular nucleus (EP) converges onto ventromedial (VM) and ventro-antero-lateral (VAL) thalamic nuclei (Guo et al., 2018; Sakai et al., 1998). GABAergic basal ganglia neurons have high tonic firing rates in vivo (Delong, 1971; Lobb and Jaeger, 2015; Ruskin et al., 2002), resulting in tonic thalamic inhibition through powerful GABA-A synapses (Bodor et al., 2008). Therefore, the motor thalamus is a key structure at the cross road between the BG-thalamo-cortical (Bosch-Bouju et al., 2013) and cortico-thalamic loops (Guo et al., 2017; Theyel et al., 2010). Notably, the motor thalamus in rodents is entirely made up of glutamatergic neurons projecting to cortex (Bosch-Bouju et al., 2013).

Recent studies in mice show that the VM thalamic nucleus forms a closed loop with anterolateral premotor cortex (ALM) and that this loop is required for persistent activation of neural activity related to movement preparation and decision making during the delay period of a cued left/right lick task (Guo et al., 2017). However, the processes by which the basal ganglia interact with this cortico-thalamic motor planning loop and may influence specific aspects of decision making remain unknown.

To address this important question on how the basal ganglia may influence motor control we recorded neural activity in VM/VAL with 4-shank silicon probes, while optogenetically activating BG terminals in this structure during distinct periods of task performance. In order to build on previous work, we adopted a version of the delayed left/right lick task used previously to discern thalamo-cortical dynamics in decision making and movement initiation (Guo et al., 2014a, 2017). We found that stimulating BG GABA release in VM/VAL just before the GO signal at the nigro-thalamic synapse lead to a decrease in impulsive licks (defined as premature action resulting in task failure) as well as an increase in the amount of omissions (defined as licking too late or not at all). Detailed analysis of neuronal activity revealed that the optogenetic stimulation produced spike rate changes in VM thalamus that were very similar to the signature of spontaneously occurring omission trials. Together our results indicate that this specific pathway contributes to controlling the proper timing of action release.

## RESULTS

### Optogenetically targeting a specific nigro-thalamic pathway

In order to target the nigro-thalamic pathway (Figure 1A) we used transgenic Vgat-ires-Cre knock-in mice (Vong et al., 2011) that express the Cre-recombinase under a Vesicular GABA Transporter (VGAT) promoter, therefore targeting GABA releasing neurons, including GABAergic SNr neurons. The SNr of VGAT-Cre mice was injected with an adeno associated viral vector (rAAV2) containing a Cre-dependent Channel Rhodopsin 2 (EF1a-DIO-hChR2(E123T/T159C)-EYFP). SNr terminals expressing ChR2-EYFP in the motor thalamus were activated using a blue laser (473 nm) passing through an optic-fiber (200µm diameter, 0.22 NA). Electrophysiological signals from neurons in VM/VAL thalamic nuclei were recorded using 4-shank 32-channel silicon probes (Neuronexus, A4×8-7mm-100-200-177). Both electrode placement and injection site (Figure 1A) were verified histologically at the end of the experiment (Figure 1B). We verified that the EYFP was successfully expressed in both the SNr and VM/VAL using immunohistochemistry and fluorescence confocal microscopy (Figure 1B). Electrode placement was matched to the Allen Institute brain atlas (Shamash et al., 2018), allowing us to define VM/VAL recording sites (Figures 1C). Finally, we used a photo-tagging strategy to further confirm electrode placement and identify which neurons responded to SNr photo-stimulation (Figures 1C, S1A, S1B and S1C). To do so, blue light pulses (1.5s pulse every 3sec during 2-3min) were applied in the VM/VAL recording area to activate SNr terminals, with an expected direct inhibition of thalamic neuronal activity (Figure 1D). A post-hoc statistical analysis (Wilcoxon test, nstim = 54 +/- 30 laser pulses per session) was used to determine which neurons had their firing rate significantly modified by photo stimulation (Figure 1C). We defined 3 photo-tagged populations: 1- the neurons photo-inhibited (SvTh-; most likely inhibited directly by SNr input), 2-neurons photo-excited (SvTh+; likely indirectly excited via disinhibition, see discussion), 3-neurons with activity not significantly altered by the light pulses (NOL). The comparison of the spatial distribution between the three populations within the thalamic recording area indicated overlapping locations (Figure S1A, S1B and S1C), except along the Medio-Lateral axis between SvTh- and SvTh+ and along the Antero-Posterior axis between SvTh- and NOL (Figure S1A).

**Figure 1.**
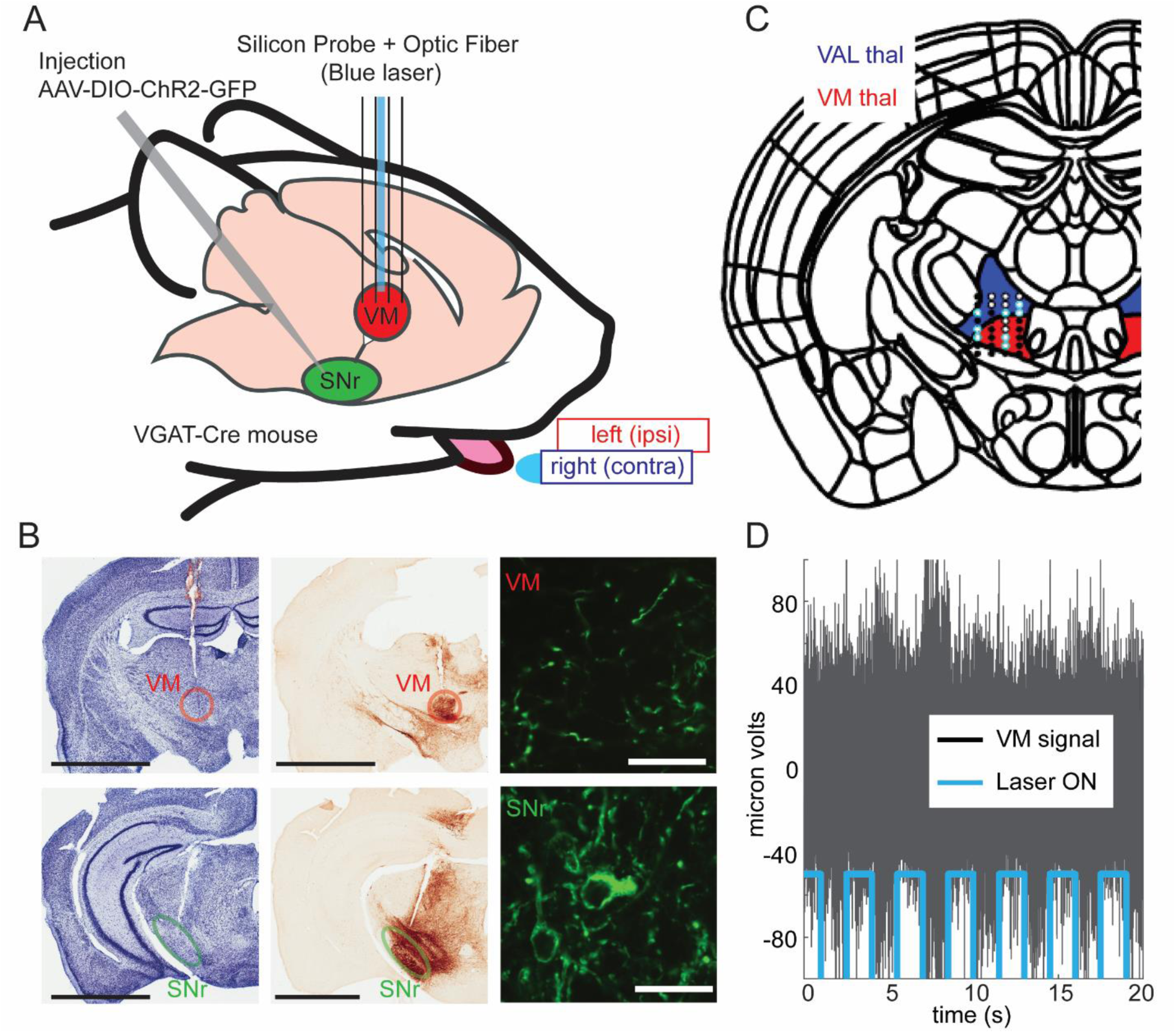
An experimental design combining electrophysiology and optogenetic to target a specific nigro-thalamic pathway. A. Schematic of the experimental paradigm. Slc32a transgenic mice were injected with AAV-DIO-ChR2-EYFP in the SNr to express ChR2. Single unit activity in VM and VAL thalamus was recorded using silicon probes. SNr terminals were optogenetically activated by illumination with blue light through an optic fiber attached to the silicon probe. Neurons were recorded in head-fixed mice performing a cued left/right licking task. B. Histology was performed to locate the position of the electrodes (top row) and injection sites (bottom row). Nissl staining (left column), immunohistochemistry with anti-GFP revealed with horseradish peroxidase (middle column), and confocal microscopy (right column). Notice in those representative examples that the fluorescence (green) indicate that GFP was expressed in the soma of SNr neurons (bottom right panel) but not in the soma of VM neurons, as GFP in the VM area was only expressed in GABAergic SNr terminals (top right panel). Scale bars: black = 2mm; white = 20µm. C. Example of electrode track reconstruction in VM (red area) and VAL (blue area). Channels excluded for analysis (black dots), analyzed VAL channels (purple dots) analyzed VM channels (green dots), channels with optogenetically responsive cells (yellow rings). D. Example of a channel in VM for which the photo-stimulation (blue 473 nm laser, 10mW, 1.5 sec ON, then 1.5 sec OFF) significantly reduced the multi-unit activity recorded in-vivo (Wilcoxon test; P<0.00001).

These results validate that our experimental paradigm allows us to specifically record the electrophysiological activity of neurons from VM/VAL motor thalamus where the laser stimulates specifically the GABAergic terminals projecting from SNr.

### Impulsive and omission trials remain after successful learning of a bi-directional delayed licking decision task

In order to better understand the role of the nigro-thalamic pathway in decision making, we used a licking task in which mice had to make a binary choice based on a sensorial discrimination and with a delay separating perception and action. Similar tasks have been used and validated by previous studies in head fixed mice (Guo et al., 2014ab, 2017; Li et al., 2015). In our version of the task (Figure 2A and supplementary video), the mice received a mild air puff (750ms duration) on left or right whiskers and withheld their response during the delay period (750ms duration). A sound Go-cue indicated the end of the delay and the start of the response period (1500ms duration). During the response period the mouse had to indicate that they remembered the side of the previous air-puff stimulation by licking into the lick-port located on the same side. A water reward was obtained, only if the first lick was to the correct side and not later than 1.5 seconds after the Go cue. An inter-trial-interval (ITI) of a minimum of 3 seconds was given between trials and the next trial started only after the mice stopped licking for 2 seconds (Figure 2A). The mice could succeed (correct trials, ‘cor’) or commit 3 types of errors (Figure 2B), lick too early (impulsive trials, ‘imp’), lick too late (omission trials, ‘omi’), or lick on the wrong side (error side trials, ‘eSi’). Mice were trained for 6-8 weeks for an average of 25+/-3 sessions, until they reached a 70% criterion performance for licking the correct side. After training, 5 mice were successfully recorded during the task across 2060 trials in 15 sessions (137+/-40 trials per session). In these trials, mice were licking the correct side first in 87% of trials, while making 13% +/-10% of side errors sides (Figure 2C). Importantly, omission and impulsive trials did not vanish after training (Figure 2C). The proportion of these impulsive and omissive trials gave us an important window on assessing the timing of lick initiation with respect to optogenetic nigral terminal stimulation.

**Figure 2.**
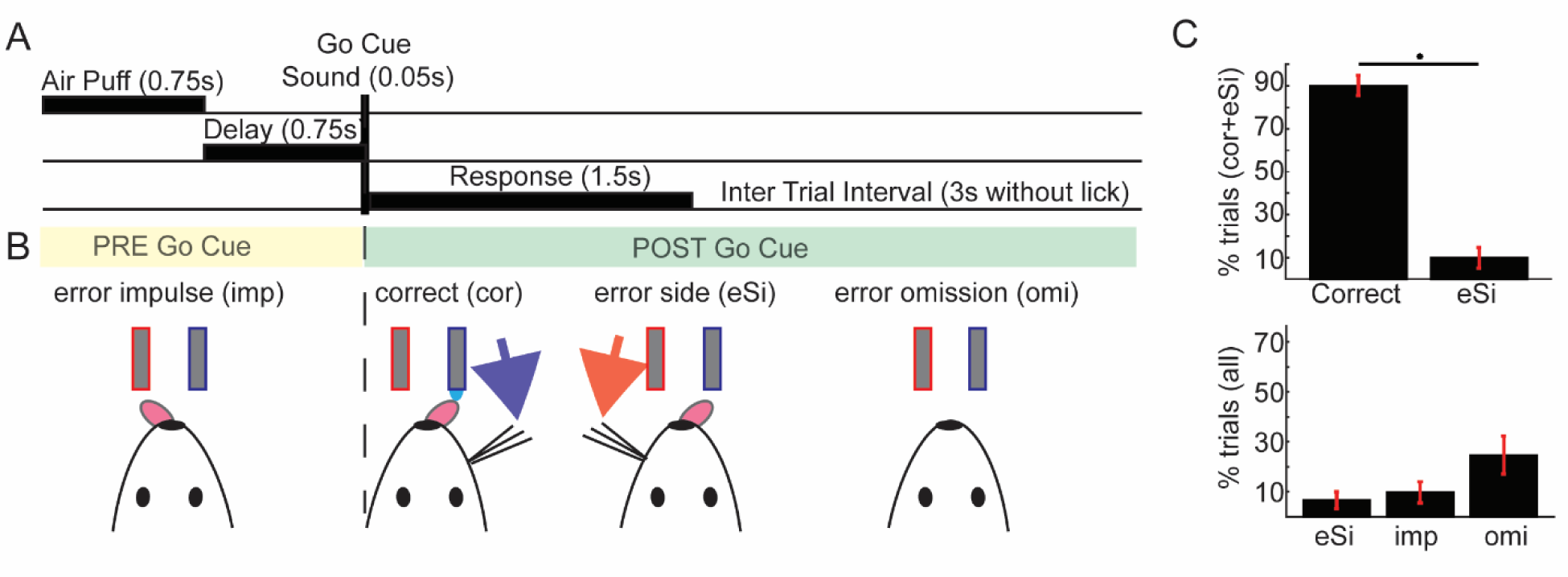
Cued choice Delayed Response Task. A. Schematic time course of a task trial. At the start of each trial, a mild air puff (sample period) was delivered pseudo-randomly on the left or on the right whiskers of the head fixed mouse. After the sample period, the mouse had to withhold licking during the delay period. The end of the delay was indicated by a Go signal after which the mouse could initiate the response lick. If the mouse first licked the correct spout during the response period, a water reward was delivered. Trials were separated by a 3 s minimum intertrial interval (ITI). Any licks during the ITI lead to a restart of the counter to starting the next trial to 2 s, ensuring that at the onset of each trial the mice had completely stopped licking from the previous trial. B. Schematic of the four possible trial outcomes: 1-Error impulsive lick (before Go cue), 2-Correct lick 3-Error lick wrong side, 4-Error no lick (omission). C. Bar graph (mean and SEM) of the behavioral results (n=15 sessions, 5 mice, 2060 trials) obtained after training periods, during electrophysiological recording. Top panel: Two sample ttest (n=15), ***P<0.0001, with 87.4% +/- 10.4% correct trials (mean +/- SD). Bottom panel: One sample ttest (n=15), **PeSi < 0.005; ***Pimp and Pomi < 0.00005. The amount of error related to timing of action (omi + imp; 43.2%+/- 14.7%) were significantly higher than the amount of errors related to directionality (eSi; 8.9% +/- 8.1%), two sample ttest (n=15), ***P<0.00005

VM/VAL thalamic neurons displayed task related firing rate changes that were sequentially organized but non-homogeneously distributed in time.

To determine the relation of VM/VAL activity to our delayed response task, we recorded the activity of 462 VM/VAL thalamic neurons during the task (Figure 3A and 3B).

**Figure 3.**
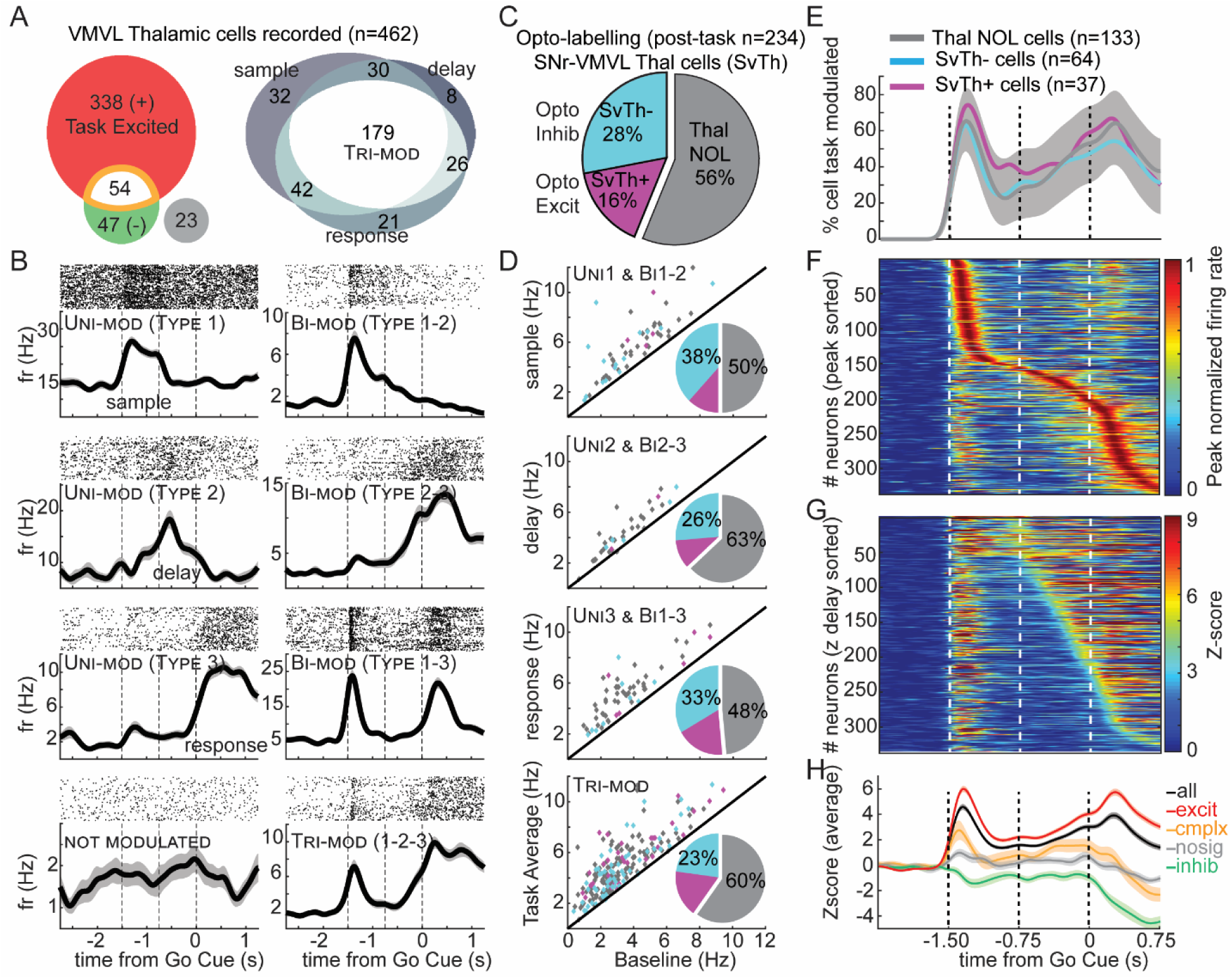
Task related neuronal responses and their distribution in time across nigral responsive and non-responsive neurons. A. Venn-diagrams. Left: VM/VAL neurons with firing rate significantly changed during the correct trials of the task (abs(z-score)>3), increased only (red), decreased only (green), both increases and decreases (orange), and not significantly modulated (grey). Right: VM/VAL neurons (Among the 338 task-excited) with firing rate significantly changed during the 3 periods of the task, sample, delay, response (z-score>3). B. Examples of raster plots and their corresponding Peri Stimulus Time Histogram (PSTH) centered on the Go cue for all correct trials. Each panel represents the spiking activity of a single unit recorded in VM/VAL during the task. Dashed lines represent the onset of task epochs. (Shaded:1 SEM.). C. Percentage of opto-tagged cells (among 234 cells tested post-task). SvTh+: optogenetically activated neurons (magenta). SvTh-: optogenetically inhibited neurons (cyan). ThalNOL: thalamic no optical response cells (grey). D. Scatter plot comparing baseline and period specific average firing rates for a specific subpopulation of cells. SvTh-(cyan), SvTh+(magenta), ThalNOL (grey). E. Percentage of cells with a significant (z-score>3) modulation in rate during the task. The shaded area represents the limit of the confidence interval (5% two-sided) resulting from a bootstrap (500 iterations, 30 cells randomly picked). 75 ms Gaussian Kernel convolution. F. Distribution of the peak normalized firing rate across time (sorted by time of the peak) for the 338 task-excited VM/VAL neurons. G. same as F but using z-score across time (sorted by first occurrence of z>3 during the delay). H. Z-score average in function of time for neuronal groups defined in A based on their firing rate properties.

First, we wanted to determine how VM/VAL individual neurons were overall modulated during the corrects trials of the task (Figure 3). We found that more than 80% of the neurons responded by increasing their firing rate to one or multiple periods of the task (Figure 3A, B). Due to the low number of inhibitory responses we decided to only further analyze the 338 neurons that exclusively increased their firing rate in relation to behavior (z-scored firing rate changes > 3). As we categorized neuronal responses by strongest-response period, we noticed that a large part of the population was task-modulated in a combination of periods (Figure 3A right and 3B right), with either 2 peaks or one peak and a ramp (bi-modal) or 2 peaks and a ramp (tri-modal). These results show that multiple time points of the task are represented at the single neuron level.

Second, we wanted to know if the neuronal population affected by optogenetic stimulation represented a distinct category of responses. Among the 234 cells optogenetically tagged post-task (Figure S1A), 28% were photo-inhibited (SvTh-), 16% photo-excited (SvTh+), and 56% none showing photo-modulation (NOL; Figure 3C). Neurons in all 3 opto-tagged categories were found in near-equal proportion to their respective totals across all response types in the task (Figure 3D). Furthermore, the distribution of significant firing rate increase (z-score >3) was not different across the 3 cell populations (Figure 3E) indicating that nigral input affect all categories of single cell response in our task.

Third, we investigated the population activity to determine the temporal distribution of rate increases in task-responsive VM/VAL neurons. The results demonstrate two distinct increased proportions of rate changes, the first shortly after whisker stimulus onset, and the second after the Go-Cue (Figure 3F). In order to see if the increased neuronal activity during the response period reflected motor activity associated with licking or preceded such licking, we sorted the activity by z-score during the delay period (Figure 3G). We observed that most neurons ramped up in spike rate before any licking, indicating that they were not solely associated with movement itself, but related to preparatory activity anticipating the action.

Finally, we computed the mean firing rate for the overall population of VM/VAL neurons (Figure 3H) and observe an overall tendency to respond to both the initiation of the trial and the initiation of the action. Note that inhibitory responses were also associated with sensory cue and response period onsets, but no inhibitory ramping was observed.

In supplementary Figure S2, we replicate previous observations (Guo et al., 2017) that VM/VAL single units presented lateralized pattern of activity during all periods of the task. We found similar proportion of ipsilateral and contralateral coding in VM/VAL during the task (Figure S4A). Figure S4B, provide representative examples of single cells with lateralized activity.

In supplementary Figure S3 we provide the stereotaxic coordinate distribution within the VM/VAL recording area of each response type in the task. We found that: 1-along the anteroposterior axis, task inhibited cells (AP = −1.361mm +/- 0.221) where statistically more posterior (Wilcoxon test, Figure S3B) compared to task activated cells (AP = −0.281 mm +/- 0.233); 2-along the mediolateral axis ‘delay activated neurons’ (ML = −0.787mm +/- 0.103) were statistically more medial (Figure S3D) compared to neurons with other response types (ML = −0.902mm +/- 0.190) and ‘stimulus activated neurons’ (ML = - 0.992mm +/- 0.200); 3-along the dorsoventral axis, neurons with unimodal responses (DV = −4.008mm +/- 0.230) where statistically more ventral (Figure S3F) than neurons with multimodal responses (DV = −3.914 mm +/- 0.221, Figure S3F). These differences in mean location suggest that a spatial functional architecture within VA/VML exists, with however largely overlapping populations with different response types.

### Optogenetic stimulation of SNr terminals in VM/VAL reduces impulsive trials and increases omission trials

The results from the previous sections showed a ramping of activity before Go-Cue presentation, which may be related to preparation of movement initiation.

We hypothesized that higher levels of ramping activity may be associated with earlier movement onset (impulsive movement), and low levels with a failure to move (lick omissions). In order to test this hypothesis, we stimulated the SNr GABAergic terminals during the delay period of the task (20% of the trials, randomly selected; Figure 4A). If there is a link between VM/VAL activity and the likelihood of lick initiation, activation of this inhibitory input should reduce the amount of impulsive trials and increase omissions. We indeed found a significant reduction in the amount of impulsive trials during opto-stimulated trials compared to non-opto trials as well as a significant increase in omission trials (optoON vs optoOFF, two tailed paired ttest for 11 sessions, 4 mice, Pcor = 0.056, PeSi = 0.076, Pimp = 0.030 Pomi=0.025, Figure 4B). Interestingly our optogenetic manipulation could not be detected by only analyzing the correct trials, indeed the increase in omissions denotes an increase in failures whereas the decrease in impulsive trials allows for an increase in correct trials. This opposite effect on the probability of a correct trial allowed for an overall statistically insignificant change in correctly executed trials (Figure 4B, ‘cor’). Note that the ratio of left (ipsilateral) and right (contralateral) trial outcomes remains statistically unchanged by the optogenetic manipulations (Figure S4D).

**Figure 4:**
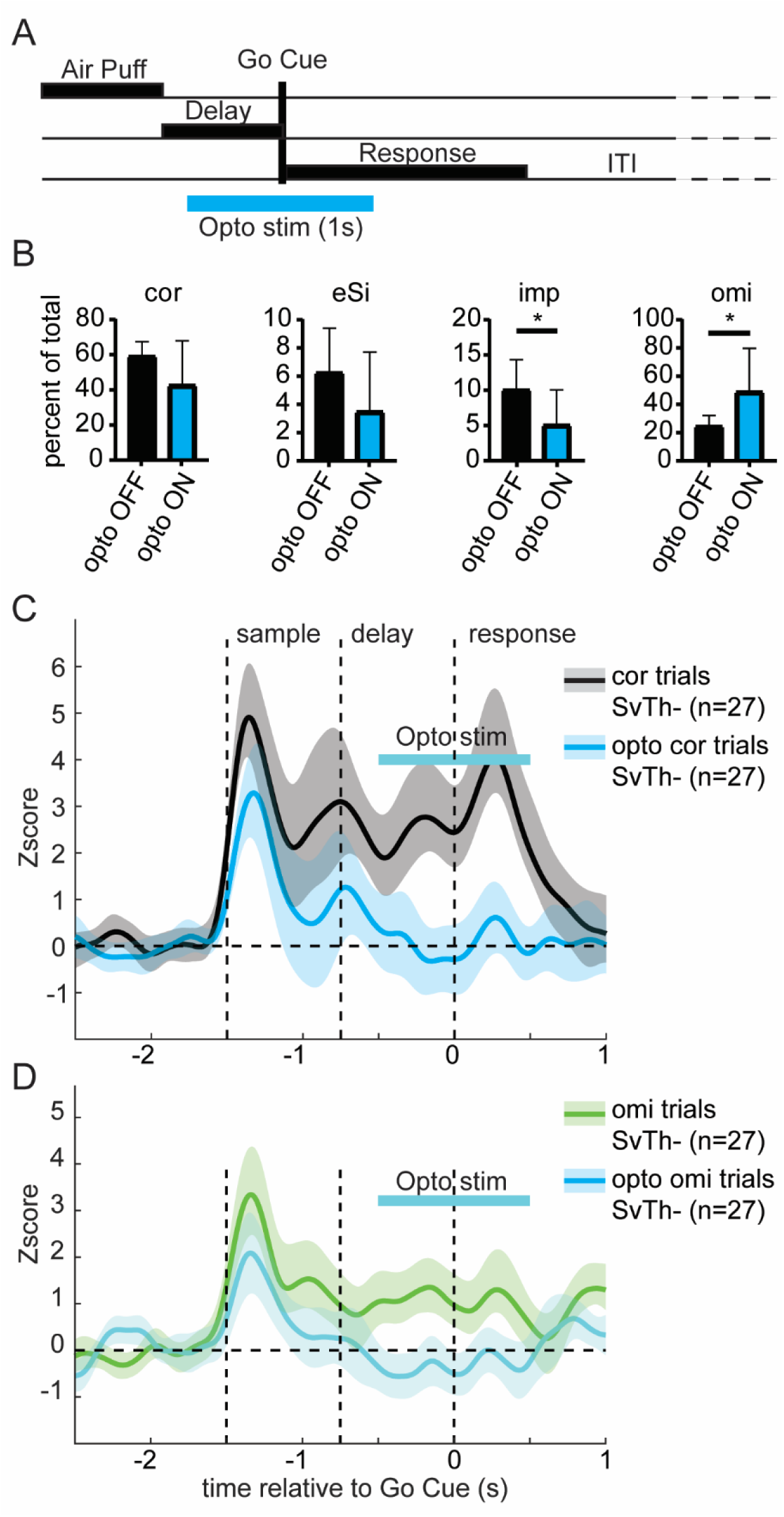
Optogenetic activation of SNr terminals in VM/VAL reduced premature licking and increased omissions. A. Schematic time course of a trial with optogenetic stimulation (blue line). Stimulation occurred in 20% of the trials (selected in a pseudorandom manner). A blue laser was turned on continuously for 1 second during the delay period 500 ms before the Go cue. B. Behavioral results with photostimulation: We tested the effect of the photostimulation of the occurrence of each trial outcome type (two tailed paired ttest). The photostimulation reduced the occurrence of impulsive trials and increased the occurrence of omission trials. (4 mice, 11 sessions) C. nigro-thalamic population (SvTh-n=27) averaged z-scored firing rate across correct trials with or without photostimulation (Opto stim). Only session with at least 7 correct opto trials were used (3 mice, 5 sessions). The neuronal activity is strongly suppressed during the delay period and subsequent response periods while the laser is ON (blue curve) compared to when the laser is OFF (black curve). Shaded: 2 SEM. D. Same as C, but for omission trials.

We then assessed the effect of the optogenetic manipulation during the task on the population activity for the VM/VAL neurons (n=89) that were previously photo-tagged outside of the task. The photo-inhibited neurons (‘SvTh-‘, n=27) presented a clear reduction in firing rate specific to the opto-stimulation epoch during the delay period of both the correct trials (Figure 4C) and omission trials of the task (Figure 4D). The non-photo-modulated neurons (‘NOL‘, n=53) showed a lesser reduction of firing rate, therefore contributing to the decrease of ramping that explains the changes in behavior (Figure S4A). The photo-activated neurons (SvTh+ n=9) did not display photomodulation during the task (Figure S4B). The grand average of all neurons tested (n=89) overall showed a robust overall decrease firing in rate during the delay and response periods (Figure S4C). In addition, we analyzed the SvTh-population average firing rate change separately for contralateral and ipsilateral trials with or without opto-stimulation. We found that in both cases the firing rate was reduced to baseline during the stimulation, indicating that the photo-inhibition was not specific for left or right trials (Figure S5E and S5F).

These results show that the opto-stimulation prevented the nigro-thalamic neurons from ramping up during the delay epoch, in agreement with our hypothesis that such activity may induce impulsive licking, and the lack of this activity induce omissions. These results indicate that during movement preparation, the level of activity in VM/VAL modulates the timing of lick initiation.

### VM/VAL neuronal activity reflected the latency of action initiation during spontaneous impulsive and omission trials

We next hypothesized that a direct correlation exists between neural activity levels in VM/VAL during the delay period and the timing of lick initiation in the absence of optogenetic stimulation as well. In order to address this question, we compared activity rates during correct trials versus early impulsive lick (imp) or no-lick trials (omi).

To define neurons selective for impulsive and omission trials, we computed z-scored mean rate differences for each neuron between different trial outcomes: dZomi = Zomi-Zcor and dZimp = Zimp - Zcor. Neurons were categorized as omission responsive if dZomi < −3 and impulsive responsive if dZimp > 3. Among 338 VM/VAL task responsive neurons, we found that 188 had their firing rate significantly modulated during either lick omission (170 cells), or impulsive lick initiation (60 cells). Notice that 42 cells among the imp responsive cells were also omi responsive, (labelled as both in Figure 5A). The PSTH histograms of 3 representative neurons with both omi and imp selectivity are shown in Figure 5B. These PSHTs clearly support our hypothesis at the single cell level, namely that the firing rate was reduced significantly during omission trials (green) and was increased during impulsive trials (violet).

**Figure 5:**
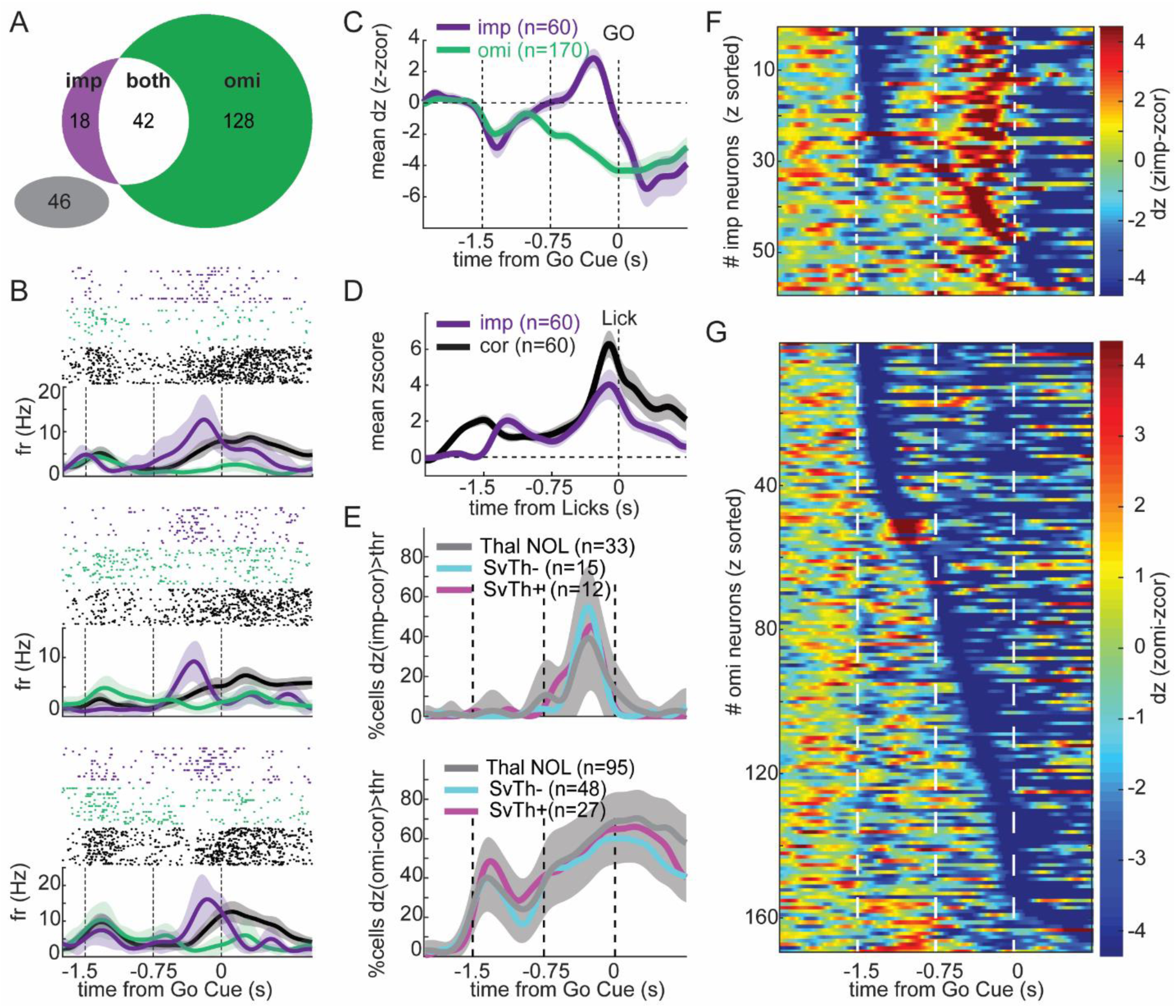
VM/VAL neuronal activity during delay is reduced during omission and increased before premature licking. A. Venn Diagram of 234 VM/VAL task-excited neurons with a firing rate difference during the delay epoch (dZ>3) either between cor trials and omi trials (green), imp trials (violet), both omi and imp (white) or no difference (Grey). B. Example of single unit activity separated by trials type, impulsive (violet), omission (green) and correct (black). Each panel represent a raster plot and a PSTH for each type of trials for an example VM/VAL neuron. C. Population average change in normalized firing rate during imp and omi trials relative to correct trial during the delay epoch. The horizontal dashed lines at zero dZ indicate no difference to correct trials. Notice how during the delay the two population diverge from zero on opposite side indicating that during delay impulsive responsive neurons tend to over-activate while omi neurons tends to hypoactivate. D. Mean z-score in function of time, align to the lick, between cor and imp trials. Both ramp up before the lick. E. Average distribution of activity difference between correct and impulsive trials (dZ>3; top panel) or omission trials (dZ<3; bottom panel) for each imp and omi responsive neurons respectively subdivided into 3 populations SvTh- (cyan; 25% of the imp cells; 28% of omi cells), SvTh+ (magenta, 20% of imp cells, 16% of omi cells) and ThalNOL (grey, 55% of imp and 55% of omi cells). The temporal distributions of their modulation were not significantly different across groups (shaded grey area represent the bootstrapped i.e. obtained by averaging over 1000 iterations with 10 cells picked randomly among all group at each iteration). F. Cell by cell distribution of activity difference between correct and impulsive trials (dz=Zimp-Zcor; top panel) or omission trials (dz=Zomi-Zcor; bottom panel) for each imp and omi responsive neurons respectively sorted by dz value. Each line is one neuron.

To determine the timing of imp and omi selectivity, we plotted the population average for each group of cells (Figure 5C). Omi selective neurons (green line) on average showed a gradual decrease of activity during omi trials that started even before the offset of the whisker air-puff and reached a peak mean difference of 4 standard deviations at the time of Go-Cue presentation. Imp selective neurons (purple line) showed a steep increase in activity only towards the second half of the delay period. Because the impulsive trials by definition contain a lick during the delay, we checked for the possibility that imp specific activity was just corresponding to lick motor activity by plotting the z-scored population firing rate aligned to the lick instead of the Go-Cue (Figure 5D). We found that firing rate ramped up before the first lick in both imp and cor trials, though it ultimately reached a higher level just before lick initiation in trials were licking was correctly executed after the Go-Cue. Interestingly, both omi and imp selective neurons showed a decrease in activity at the onset of the air-puff stimulus for omi and imp trials, respectively (Figure 5C). This finding suggests that even early during each trial a failure can be predicted by a lack of stimulus responsiveness in the same neurons that are later selective for failures. The timing of Omi and imp selectivity was equally represented across the three photo-tagged groups with different response types to SNr terminal activation (Figure 5E), indicating that it was not a specialized property for neurons with a particular response to SNr input. Note that the proportion of cells with selectivity was also similar across all groups, namely between 23 and 32% for imp selectivity and 72 and 73 % for omi selectivity (see Figure 3C for group totals).

Finally, in order to resolve the timing of imp and omi selectivity at the single neuron level, we plotted the z-scored rate difference for each neuron as a heat map. For imp selective neurons during imp trials, a consistent strong increase in spike rate was present during the second half of the delay period, while about 50% of these neurons also showed a strong decrease of firing during the stimulus period (Figure 5F). Omi selective neurons during omi trials showed a reduction in activity that could start at varying time points beginning with the stimulus onset, and generally lasted beyond Go-Cue presentation. We also examined the spatial distribution of imp and omi selective neurons but found no significant differences in spatial distribution (Figure S5).

### Impulsive and omission behavior can be predicted by the activity of VM/VAL neuronal assemblies receiving input from the SNr

Our previous analysis of VM/VAL neural activity revealed ramping activity prior to lick onset. We hypothesized that modulation of this activity for different trial outcomes contained sufficient information in order to predict each outcome. To test this hypothesis, we trained a logistic classifier to predict trial outcome (e.g. omi vs cor) based on the neuronal activity of randomly chosen subpopulations of neurons. To further address the question if such prediction was limited to neurons responsive to SNr terminal activation, we analyzed SvTh- and NOL neurons separately.

We found that the classifier was indeed able to predict trial outcomes for omission versus correct trials (Figure 6A), impulsive versus correct trials (Figure 6B), and omission versus impulsive (Figure S6). In all cases, we show that the classifier performance was better than chance (dashed line) and improved as a function of the number of neurons used to train it. We found no difference when comparing between neurons from SvTh- (cyan) and NOL (black), indicating that both populations contained information to predict trial outcomes outcome.

**Figure 6:**
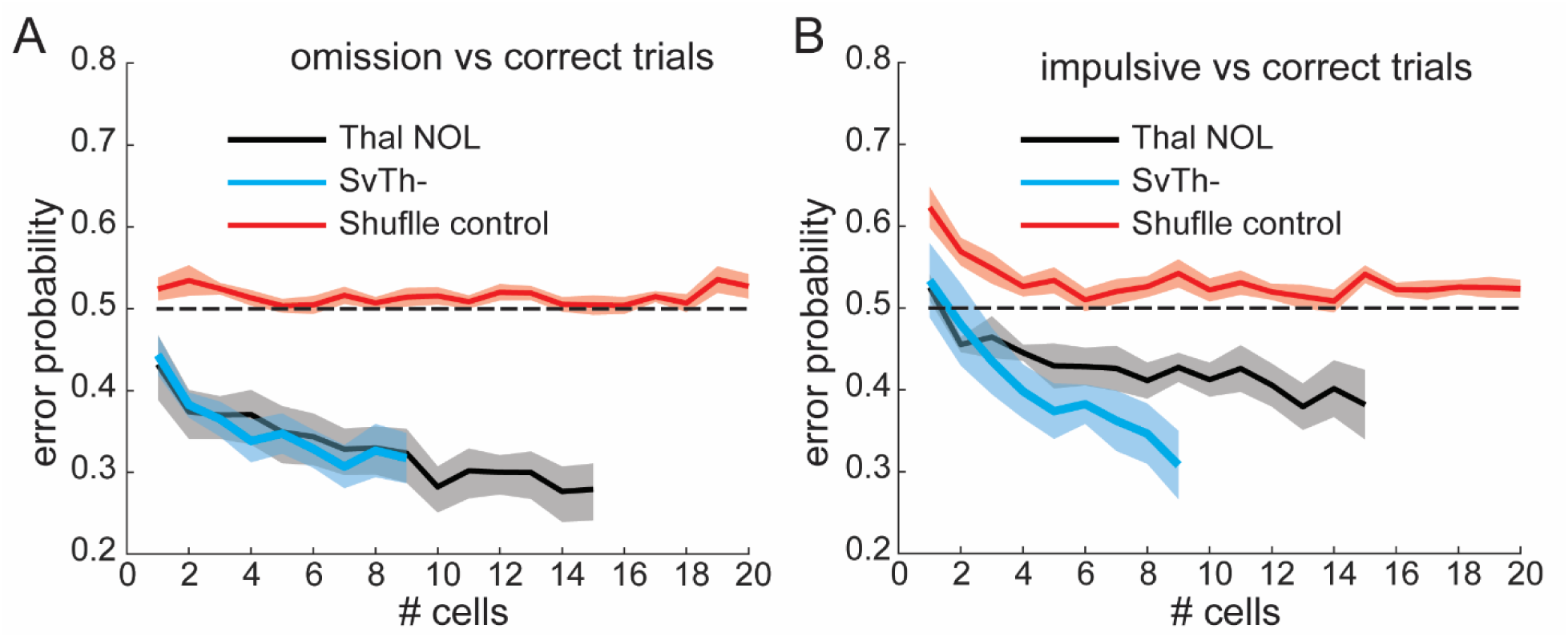
Thalamic Cell assembly encode the type of action that are prepared during the delay of the task. A. Average performance of Logistic classifier on discriminating omission vs correct trials based on the firing rate of neuronal assemblies of different sizes (randomly selected across the VM/VAL populations SvTh- or NOL) during the delay of correct trials only (no licking during delay). 10 cross validation. 1000 iterations for each group size. Cells were randomly selected at each iteration. For a shuffle control, trial outcomes were randomly assigned to each trial spike train. B. Same as A but comparing impulsive trial vs correct trials. Note that because the mouse licks during delay in imp trials this classifier uses the neuronal activity of the 750ms before the lick.

## DISCUSSION

In this study we showed that firing rates in basal ganglia input recipient motor thalamic nuclei (VM/VAL) in a task with a delayed Go-Cue predict the timing of response licks to determine correct trial outcomes, or error trials with impulsive early licks or lick omission. On the population level, omission trials were associated with a clear firing rate reduction during the delay period between sensory and Go-Cue, whereas impulsive lick trials showed a pronounced increase in firing rate over correctly executed trials. Single neurons showed highly significant rate increases during several distinct periods of the behavioral task, often peaking shortly after trial onset and again after the Go-Cue. Optogenetic activation of SNr GABAergic terminals in VM/VAL resulted in an increase of omissions and decrease in impulsive licking. Spontaneously occurring impulsive trials could be predicted by a linear decoder based on enhanced firing rate increase during the delay period, which could reflect a natural over-motivation for the reward. Vice versa, spontaneously occurring lick omissions could be predicted based on firing rate decreases in task related responses.

When optogenetically stimulating GABAergic terminals in VA/VAL thalamus outside of the task we observed the expected decrease in thalamic neural firing in a proportion of 28% of neurons, while 56% showed no significant spike rate change and 16% showed an increase. However, no functional separation in terms of task responses was observed between these populations, and the non-responsive neurons on average did show a firing rate decrease when stimulated during the delay period in the task. It seems therefore likely that nominally unresponsive neurons did not have sufficient ChR2 expression in the stimulated SNr terminals rather than not receiving nigral input. Neurons that showed increases with stimulation outside of the task were most likely not directly excited by GABAergic nigral input but received indirect excitation through network interactions. The most likely candidate for these network effects in motor thalamus is presented by the reticular nucleus of thalamus (RTN), which is excited by thalamocortical (TC) neurons, and in turn inhibits these neurons (Ando et al., 1995). Our optogenetically inhibited TC neurons therefore are likely to lead to reduced RTN excitation, which in turn would broadly disinhibit TC neurons that are not themselves inhibited by the SNr. Nevertheless, we observed a decrease in the total average activity of all neurons when stimulated during the delay period, supporting the interpretation of the behavioral outcome of increased lick omissions as a failure of increased nigral activity to trigger movement initiation.

Taken together, our electrophysiological and optogenetic results lead us to propose a functional model of basal ganglia input to VM/VAL, in which ramping activity leading to movement initiation is dampened by inhibitory SNr input (Figure 7). In this model, movement will be initiated at the time at which a threshold in VM/VAL population activity is reached (Figure 7, correct trials). If this ramp-up reaches the threshold too early it will trigger a premature action (Figure 7, impulsive trials), finally if it does never reach the threshold the prepared action will not occur (Figure 7, omission trials). Our model also explains how the opto-activation of nigral terminals in VM/VAL helps to substantially reduce impulsive trials and increased the occurrence of omission trials. This model overall generates a timing function for basal ganglia input to motor thalamus. Because basal ganglia output neurons have a high spontaneous firing rate (Delong, 1971; Lobb and Jaeger, 2015; Ruskin et al., 2002), the natural mode of basal ganglia action is to prevent premature movement initiation by dampening a thalamo-cortical feedback loop previously identified between VM/VAL and ALM cortex in the control of cued licking (Guo et al., 2017). This function is consistent with previous work showing that SNr neurons decrease in activity for action initiation, and such decreases are countermanded by subthalamic excitation when an action needs to be canceled (Schmidt et al., 2013).

**Figure 7:**
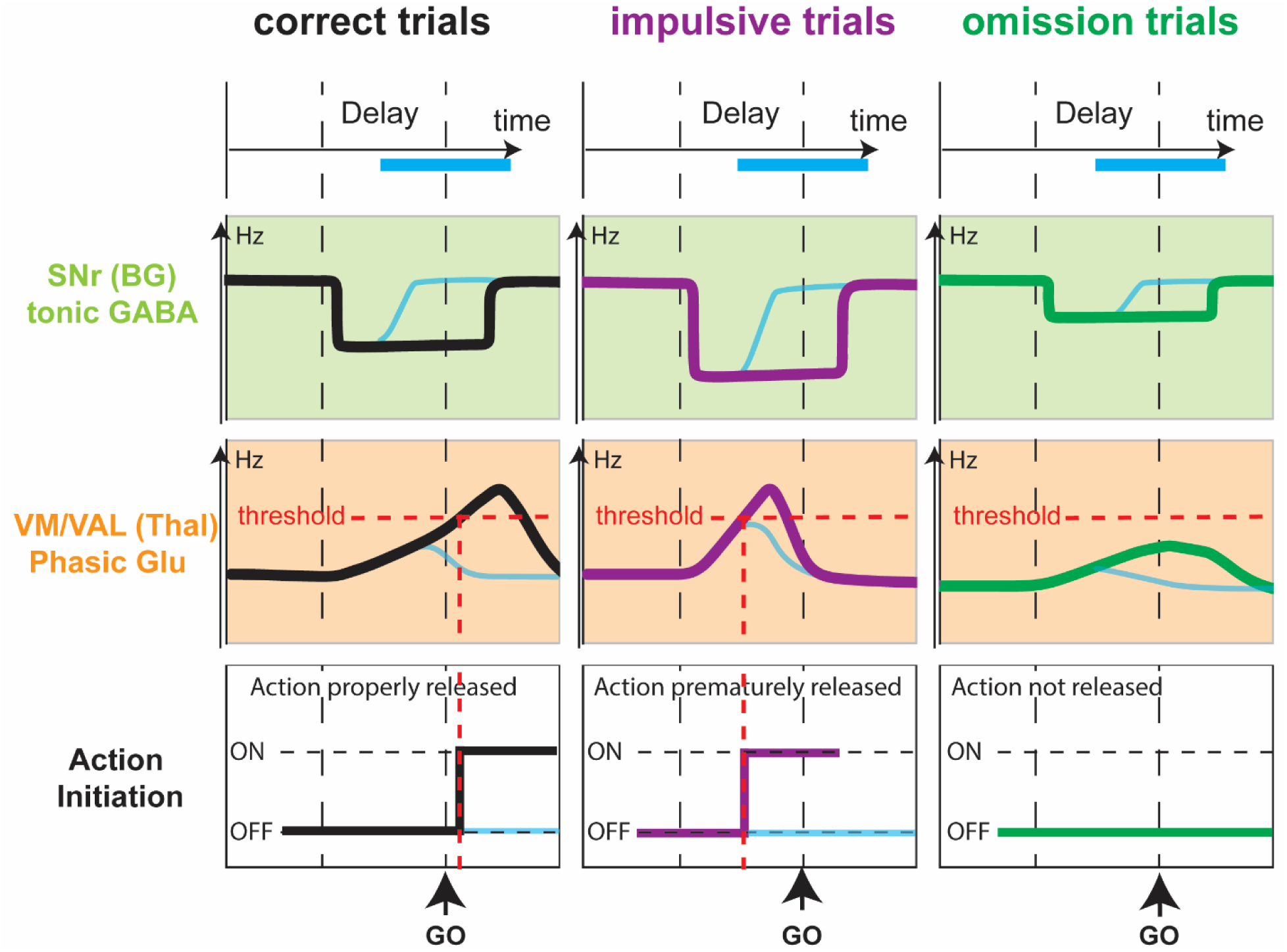
A model of nigro-thalamic activity threshold for timing of action initiation. (Left column) During correct trials, the SNr would stop his tonic firing during the delay before the GO-cue (green) leading to a ramp up activity in the receiving part of the motor thalamus (orange) that when cross a threshold leads to the downstream neural pathway that initiate the release of the action, within a proper timing windows (white). (Middle Column) During Impulsive trials, the SNr would stop his tonic firing early during the delay (green), this, combined with potential other input, will lead the thalamic activity to ramp up too fast and reach the threshold before the Go cue (orange), triggering the downstream neural pathway for initiating the action too early and therefore releasing the action during the delay instead of the response (white). (Right column) During omission trials, the SNr would not stop his tonic firing during the delay (green). The thalamic activity will not ramp up and not reach threshold (orange) The action will not be release (white). Light blue curves represent what happen during the trials with optogenetic stimulation.

Overall, our data support a model according to which the basal ganglia input to a thalamo-cortical feedback loop is responsible for the timing of action initiation in learned reward-based tasks. While our work focused on the final output of the basal ganglia to motor thalamus, previous work has shown similar ramping activity in relation to movement initiation in striatal during cued preparatory delays (Jaeger et al., 1994; Mello et al., 2015) or prior to self-initiated movements (Lee and Assad, 2003), as well as in pallidal neurons (Jaeger et al., 1994). These activities likely lead a reduction of SNr activity to release movement initiation, while in contrast subthalamic excitation of the nigra may favor stopping of movement initiation (Schmidt et al., 2013). SNr activity, in turn, is likely controlled by striatal input from the direct pathway. Indeed, optogenetic inhibition of the direct pathway (disinhibiting SNr) leads to a delay in the time of motor execution (Tecuapetla et al., 2016) in agreement with our timing model.

The vigor hypothesis posits that BG output is controlling movement vigor, as measured by an impact of basal ganglia output on action speed (Desmurget and Turner, 2008; Turner and Desmurget, 2010). This effect has also been referred to as a control of ‘movement gain’ (Anderson and Horak, 1985), which may be related to motivation (Catanese and van der Meer, 2013; McGinty et al., 2013) and cost of action (Shadmehr and Krakauer, 2008; Turner and Desmurget, 2010). Clinically, movement gain is diminished in hypometric basal ganglia disorders including Parkinson’s disease, often leading to bradykinesia and hypometria (Desmurget et al., 2004).

In our task, pre-mature actions (impulsive trials) could be seen as over-vigorous while lack of action (omission trials) can be seen as lacking vigor. We found that reduction of the BG recipient thalamic activity increase the amount of omissions and reduces the amount of impulsive trials, which is compatible with the idea of a reduction of vigor/motivation transmitted by an over-activation of the BG output leading to a dysregulation of the VM/VAL population activity as shown in Figure 7. Notably, rate increases in VM/VAL neurons in our data often lasted through the response period, and therefore our data are also compatible with a correlation between population firing rate and the vigor of the action. However, in contrast to reaching movements in primates, the speed of licking is most likely controlled by a brainstem central pattern generator for licking (Travers et al., 1997).

Our threshold hypothesis suggests that the level of neuronal activity in VM/VAL once past a certain threshold will trigger the actions even though the action was supposed to happen later, as if the motivation/vigor level went so high that the action become irresistible. On the contrary if the level of activity never reaches a certain threshold then the action will not be release at all. Therefore, in the frame of the threshold hypothesis model presented in Figure 7, the activity level in VM/VAL could be a function of a gain/vigor/motivation for a given action. This gain function combined with an adjustable threshold values, could serve as a timer for action release, as well as a system for selection based on activity level. Indeed it is possible that other actions compete (Boraud et al., 2018; Leblois et al., 2006) for the highest motivation value, and that a winner take all mechanism is implemented. Such a possibility is also compatible with our proposed mechanism (Figure 7), in which the first action to reach the threshold would be released.

## METHODS

### Mice

Data were collected from adult male and female mice between 3-6 month of age. All mice were from a slc32a transgenic line expressing Cre under the VGAT promoter in GABAergic neurons (JAX stock #028862). Animals were maintained on a C57BL/6 background and kept on a 12:12 reverse light/dark cycle. During the behavioral training periods, animals were water restricted to 85% of their body weight. Food was available ad libitum at all times.

### Stereotaxic Surgeries

Mice were anesthetized with isoflurane (4% for induction and then maintained at 2%). The analgesic Buprenorphine SR-LAB (1 mg/kg) was administered (s.c.) pre-operatively. The skull was shaved. Mice were placed on stereotaxic apparatus (Kopf). Ophthalmic ointment was applied to the eyes. Body temperature was monitored and maintained at 37.2 Celsius at all time by using a heat pad. During the procedure, sterile saline (0.1 ml s.c.) was injected as needed to maintain hydration. The level of anesthesia was monitored every 20min (toe pinch and breathing ∼1Hz). A circular scalp incision revealed the skull which was cleaned and dried. A pinhole craniotomy was drilled above the SNr (AP: −3.4 +/- 0.5mm ML: −1.4 +/- 0.5mm, relative to Bregma). A viral vector containing the transgene (rAVV2-EF1a-DIO-hChR2(E123T/T159C)-EYFP) was injected (200nl) into the left SNr (AP: −3.40 +/- 0.25mm ML: −1.40 +/- 0.25mm, DV: −4.40 +/-0.10mm relative to Bregma). After the injection, a custom machined aluminum head-bar was cemented onto the skull using dental adhesive resin (Metabond C&B, Parkell Products Inc). A pinhole craniotomy was made above the cerebellum area for the ground wire (Tungsten). The head-bar was secured with more dental cement and the entire skull area was protected with a thin layer of cement. Antibiotic ointment was applied around the cemented area.

A few days before acute electrophysiological data acquisition, a second surgery was performed on the mice that successfully completed the behavioral training. In this surgery, a small craniotomy (AP: −1.25 +/-0.50 mm; ML: −0.75 +/-0.50 mm relative to Bregma) was performed under deep isoflurane anesthesia, and protected with Kwik-Cast silicon sealant (WPI) outside of recording sessions.

### Behavioral Training

All tasks events were programmed using LabView (National Instruments™). The pre-training procedure (1-2 weeks) was adapted from Guo et al., 2014a. Mice were handled daily in the experimental room and progressively habituated to the head-fixation procedure. Once head-fixed and body restricted in a transparent plastic tube (Guo et al., 2014a), mice were placed in a dark behavioral box (ENV-018MD-EMS, med associates, Inc.) isolated from light, sounds, and electrical noise.

The training procedure (6-8 weeks) was performed daily with an interruption of one day per week during which mice had free access to water. The training was divided in 3 steps as follows:

- step1: Mice learned to respond to the Go-Cue sound by licking, within 1.5 seconds after the cue onset, on a single tube located in front of them to obtain a water droplet.
- step2: Mice learned the association between a mild air puff (oriented pseudo-randomly at each trial toward either the left or the right whiskers) and the reward side (left or right lick-port tube corresponding to the side of the previous air puff). An algorithm allowed to prevent for side preference, such that if a mouse exhibited a preference for executing trials to one side, the probability of trials to the other side was automatically increased to enforce correct execution of trials to both sides.
- step 3: Mice learned to withhold their response during a delay periods (increasing by step of 250ms until 750ms). Step transition occurred when 70% performance criterion was reached. In each step, errors (licking to early or wrong side) were signaled by an error sound and followed by a time penalty (6s).

### Electrophysiological data acquisition

Electrophysiological signals were obtained by acute insertion of silicon probes made of four shanks separated by 200µm with eight channels per shank separated by 100µm (A4x8-7mm-100-200-177, Neuronexus Inc). Neurons were recorded in the VM and VAL thalamus (AP: −1.25 +/-0.25 mm; ML: −0.75 +/-0.25mm; DV: −4.45 +/-0.45mm; relative to Bregma). Recordings were started 20 minutes after probe insertion to allow the electrode/brain interface to stabilize. A 200µm diameter optic fiber (0.22 NA) was mounted 200µm above the first channel on the 3rd shank. An internal reference channel was located 200µm above the 1st channel on the 2nd shank. Skull screws were cemented above the cerebellum and used as ground and external reference in case the internal reference would fail.

Data from electrodes and behavioral data from the task were both acquired simultaneously at 20KHz through an RHD2000 recording system (Intan Technologies). Electrophysiological data were notch-filtered at 60Hz. Video data (see supplementary video) of the mouse licking the tubes were acquired at 25Hz (Basler acA1300 camera), under infrared LED illumination. Recording of 100 frames was triggered at each trial start (Air-Puff TTL rising edge).

### Pre-processing and analysis of neuronal data

We used Matlab (Mathworks Inc) for all data pre-processing and data analysis. Spike extractions and spike sorting was obtained using Wave-Clust a method for spike detection (pass band filter; fmin=600, fmax=8000; detection threshold: stdmin=4, stdmax=30, filter order =4) and sorting (fmin=300, fmax=4000, filter order=2) with wavelets and superparamagnetic clustering (Quiroga, 2004, 2012). Each spike was represented by 40 points centered on the minimum value and had a 3ms minimum refractory period. After the automatized procedure, each cluster was re-inspected to prevent from potential type 1 or type 2 errors as well as to eliminate poorly defined or noisy clusters.

Peri-event time histogram and raster plot were centered on task event onset (Delay, Go-Cue, Licks).

All z-scores were obtained by subtracting the baseline average firing rate (frB) from each value of the trial averaged instantaneous firing rate (frT) and divided by the trial average firing rate standard deviation (sdT).

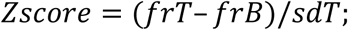

dZ are obtained by subtracting z-scores (e.g. dZ = Zomi - Zcor).

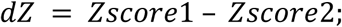

All classifiers were computed using *fitclinear* (matlab), with a Logistic learner and 10-fold cross validation. For figure 6 and S6, only sessions with at least 5 impulsive and 5 omission trials were used. Note that in Figure S6 sessions have different number of cells.

In order to compute the average, for figure 6, only sessions with at least 9 SvTh-cells recorded simultaneously were used to compute the SvTh-curves. Only sessions with at least 15 NOL cells recorded simultaneously were used to compute the NOL curves. Only sessions with at least 20 VM/VAL cells recorded simultaneously were used to compute the shuffle curves. Because for this particular analysis the number of averaged values are limited by the session with the least number of cells we arbitrarily set these number in order to maximize both the number of sessions and the number of cells in the x axis.

### Optogenetic stimulation and analysis for photo-tagging

For all photostimulation experiments, we used a blue laser (Dream Laser Inc, 473nm; 10+/-3mW at fiber end). An optic fiber patch cord (Doric Lenses; 200μm diameter; 0.22NA) was directly attached to the optrode (Neuronexus; 200μm diameter; 0.22NA) with a ceramic sleeve. Aluminum foil was carefully wrapped around the sleeve to prevent light spread to the animal’s eyes.

In-Task optogenetic stimulation consisted of 1 second long continuous pulses. Optogenetic stimuli were delivered randomly in 20% of the trials. The pulse always started 500ms before the Go-Cue during the delay period (Figure 4A).

Pre and post-task optogenetic stimuli were used for post-hoc photo-tagging (Figures 3, 4 and 6), and consisted of a series of 1.5 second pulses separated by 1.5 second inter-pulse-intervals (Figure 1D). The post-task stimulation started 1-2 minutes after the behavioral task recording was stopped without moving the optrode position. The data from the task and post-task were recorded continuously such that spike sorting was performed in one block for the entire recording. We statistically compared the spike count during 2 conditions, laser ON versus laser OFF (Wilcoxon test, nstim = 54 +/- 30 pulses per session), neurons with significant difference were considered opto-responsive. We create three categories of VM/VAL photo-tagged neurons: ‘SvTh-’ when frON<frOFF; ‘SvTh+’ when frON>frOFF and “Thal NOL” when no frON and FrOFF were not significantly different.

### Histology

At the end of the last recording session, mice were euthanized by an overdose of ketamine (i.p.), then perfused (i.c.) with phosphate buffered saline (PBS 1X) then with paraformaldehyde (PFA 4% in PBS) for tissue fixation. The brain was extracted and put in a 4% PFA solution overnight before being transferred in a 30% sucrose solution for storage at 4°C.

Brains were sliced (40µm) using a microtome and stained with 2 different methods: Nissl (Cresyl violet) and immunohistologically using anti-GFP (peroxidase). We used confocal imaging to observe GFP fluorescence on unstained slices.

### Statistical Analysis

Numerical values obtained by averaging are denoted as follows: Mean +/- SD. Statistical tests were conduct with an alpha of 5%, two samples, and two-tailed, except when specified otherwise. Statistical test results are described as significant in the text where p < 0.05. (* <0.05. **<0.005 ***<0.0005). The Wilcoxon rank sum test from Matlab is equivalent to the Mann-Whitney U test. Bonferroni correction was applied when multiple comparisons. Shaded areas in plots indicate two SEM except when specified otherwise. Specifics on the statistical methodologies and software used for various analyses are described in the corresponding sections in Results, Methods, Figure Legends, and Supplemental Figures.

## DATA AVAILABILITY

Data that support the finding of this study are available from the corresponding authors upon reasonable request.

## CODE AVAILABILITY

Codes from this study are available from the corresponding authors upon reasonable request.

## ACKNOWLEDGMENTS

We thank Dr. Adriana Galvan for helping with histological procedures. We thank Dr. Nick-Steinmetz for his help with the 3D analysis of electrode track. We thank Dr. Hidehiko Inagaki for his insightful comments on the draft. We thank members of the Jaeger lab Dr. Li Su, Dr. Arthur Morissette, Chelsea Leversedge and Jacqueline Zhu for their technical help. This work was supported by NIH 1UO1NS094302 (Jaeger, PI) and 1R01NS111470 (Jaeger, PI) under the BRAIN Initiative.

## AUTHOR CONTRIBUTIONS

JC and DJ designed experiments, JC carried out experiments, JC analyzed data, JC and DJ wrote manuscript.

